# Predicting bacterial growth conditions from mRNA and protein abundances

**DOI:** 10.1101/353433

**Authors:** Mehmet U. Caglar, Adam J. Hockenberry, Claus O. Wilke

## Abstract

Cells respond to changing nutrient availability and external stresses by altering the expression of individual genes. Condition-specific gene expression patterns may provide a promising and low-cost route to quantifying the presence of various small molecules, toxins, or species-interactions in natural environments. However, whether gene expression signatures alone can predict individual environmental growth conditions remains an open question. Here, we used machine learning to predict 16 closely-related growth conditions using 155 datasets of *E. coli* transcript and protein abundances. We show that models are able to discriminate between different environmental features with a relatively high degree of accuracy. We observed a small but significant increase in model accuracy by combining transcriptome and proteome-level data, and we show that stationary phase conditions are typically more difficult to distinguish from one another than conditions under exponential growth. Nevertheless, with sufficient training data, gene expression measurements from a single species are capable of distinguishing between environmental conditions that are separated by a single environmental variable.

## Introduction

Environmental conditions across the planet vary in terms of their capacity to support microbial life. Further, individual environments can change rapidly over time, and these changes are likely to impact the composition of microbial communities and ecosystem functions in unpredictable ways [1,2]. Microbial species composition is partially indicative of environmental conditions, particularly with regard to the presence of individual specialist species that are well adapted to unique environments [3,4]. However, many bacterial species within a community are generalists that are capable of thriving in diverse environments and must therefore sense and respond to various environmental signals [5]. For instance, *Escherichia coli* grows inside the comparatively warm, nutrient rich digestive tract of host [6] organisms but spends another portion of its life-cycle exposed to harsh environmental conditions upon being excreted and before finding another host. The mere presence of generalist species in an environment may provide little value for understanding past or current environmental conditions because their varied gene expression repertoire permits growth across varied conditions [7].

On top of their native responses to external conditions, microbial cells can be engineered to act as sensors for a variety of environmental features via rational design of synthetic genetic circuits that may, for instance, cause the cells to fluoresce upon sensing of a particular small molecule [8]. Such applications can provide a useful, low-cost diagnostic for monitoring environmental changes, but individual synthetic biology applications take time and resources to develop. Additionally, there is still a concern about releasing genetically engineered species into natural environments where they may act as low-cost sensors for pollutants or various environmental phenomena of interest [9].

To partially alleviate this concern, previous work has shown that the species composition of an environment can serve as a rapid and low-cost biosensor to indicate the presence of various contaminants according to the species abundances identified via meta-genomic sequencing [3,10,11]. However, looking at the species composition alone fails to account for the fact that gene expression patterns of individual species—particularly for generalists—may provide even higher resolution into the past and current chemical composition of environments. The extent to which gene expression patterns of individual generalist species can be used to discriminate between environmental conditions remains unknown.

Combining different ‘omics’-scale technologies is likely to provide better discriminatory capability versus only monitoring mRNA abundances, for instance, but integrating datasets is challenging due to the biases of individual methods [12] and the inevitability of batch-level effects that occur when datasets are generated across multiple labs and platforms [13,14]. These problems are further exacerbated when considering the ultimate goal of detecting different environmental conditions *in situ*.

Prior studies have looked into the question of predicting external conditions by using the cells’ internal variables [15,16]. Other studies have interrogated multi-omic datasets from different growth conditions to understand the function of regulatory networks, individual gene functions, and resource allocation strategies [7,17]. However, the main focus of many of these studies has been to understand differences in gene expression patterns across environmental conditions so as to provide insight into *internal* cellular mechanisms and pathways or to predict cellular level phenotypes such as specific growth rates. By contrast, few studies have focused on using the internal state of cells to predict external environmental conditions across a range of partially-overlapping conditions and cellular growth rates.

Here, we are interested in determining whether gene expression patterns can discriminate between environmental conditions in the absence of prior knowledge about the role and function of individual genes. Our study leverages a large dataset of transcriptomic and proteomic measurements of *E.coli* growth under multiple distinct but closely-related conditions [18]. We use mRNA and protein composition data to train machine learning models and find that highly similar environmental conditions can be discriminated with a relatively high degree of accuracy. We also investigate which conditions are more- and less-challenging to discriminate and find that prediction accuracies decrease substantially for stationary phase cells, indicating the importance of cellular growth for discriminating between conditions. Finally, we note that our accuracy remains limited by training set size such that our findings present a lower bound on the predictive power that is achievable given a greater availability of training data.

## Results

### Data structure and pipeline design

We used a previously generated dataset of whole-genome *E. coli* mRNA and protein abundances, measured under 34 different conditions [18,19]. This dataset consists of a total of 155 samples, for which mRNA abundances are available for 152 and protein abundances for 105 (Fig 1). For 102 samples, both mRNA and protein abundances are available. The 34 different experimental conditions were generated by systematically varying four parameters. Here we further simplified the experimental conditions into a total of 16, by grouping similar conditions together (e.g., 100, 200, and 300mm Na+ were all labelled as “high Na”). For the remainder of this manuscript (unless otherwise noted) we use the term “growth condition” to refer to the four-dimensional vector of categorical variables defining growth phase (exponential, stationary, late stationary), carbon source (glucose, glycerol, gluconate, lactate), Mg^2+^ concentration (low, base, high), and Na^+^ concentration (base, high). The question we set out to answer is: to what extent are machine learning models capable of discriminating between these growth parameters given only knowledge of gene expression levels, provided as mRNA abundances, protein abundances, or both?

**Figure 1:**
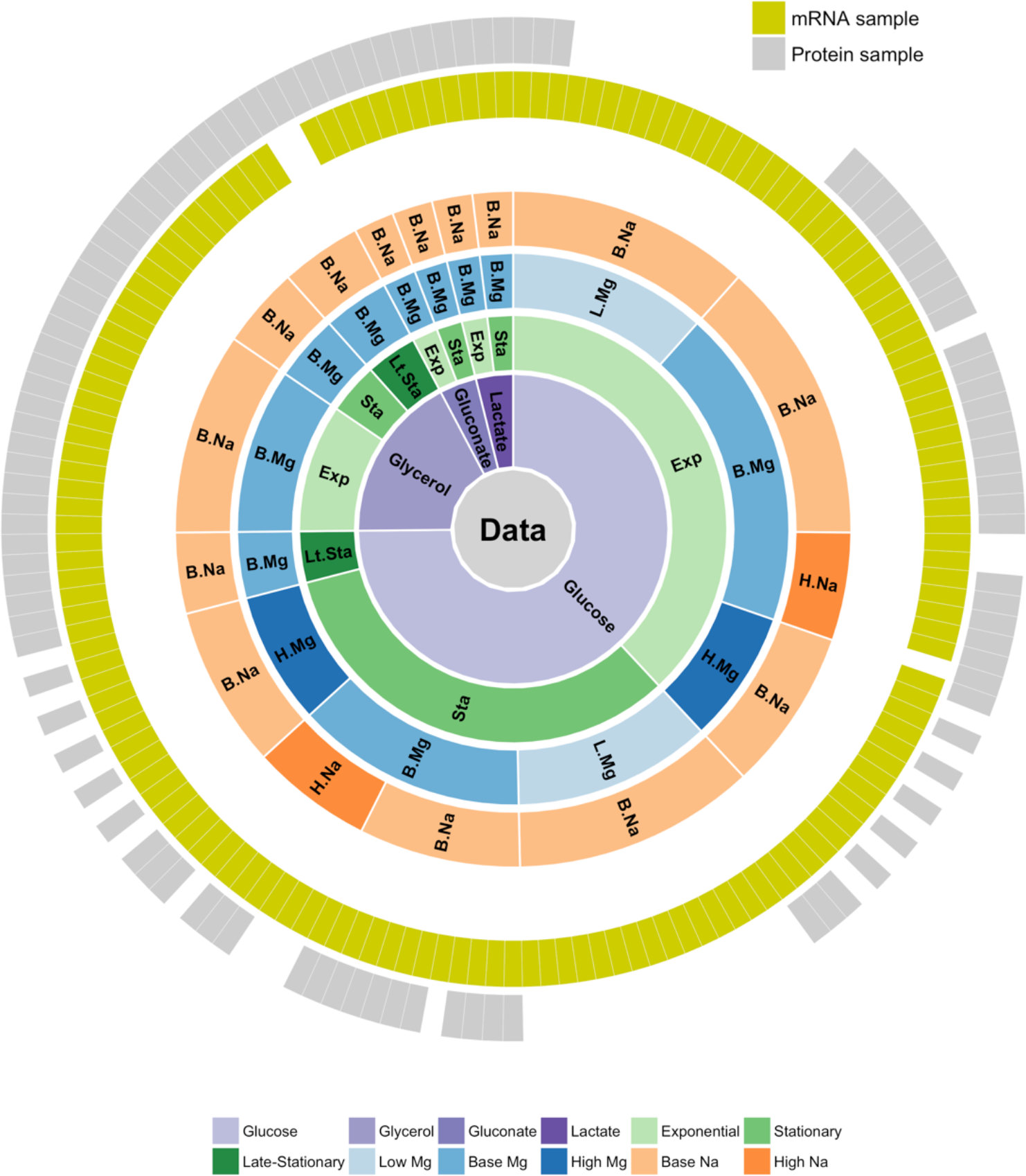
Overview of available gene expression data. Our study uses a previously published dataset consisting of 155 samples [13, 14]. 152 samples have whole-transcriptome RNA-Seq reads and 105 have mass-spec proteomics reads. 102 of the 155 samples have both mRNA and protein reads. Bacteria were grown on four different carbon sources (glucose, glycerol, gluconate, and lactate), two sodium concentrations (base and high), and three magnesium concentrations (low, base, and high). Samples were taken at multiple time points during a two-week interval, and they can be broadly subdivided into exponential phase, stationary phase, and late stationary phase samples

We applied a general cross-validation strategy and first split samples into training and test datasets. We next used the training data to fit supervised models to the gene expression data to maximize correct predictions of the labeled environmental conditions. At the training stage, we employed parameter tuning, which required a further subdivision of the training data to identify the optimal tuning parameters. Finally, we use the trained and tuned models to predict test set data and report prediction accuracy. To assess robustness of our results to the choice of training and test data, we repeated this procedure 60 times. Our pipeline is illustrated in Fig 2 and described in detail in the Materials and Methods.

**Figure 2:**
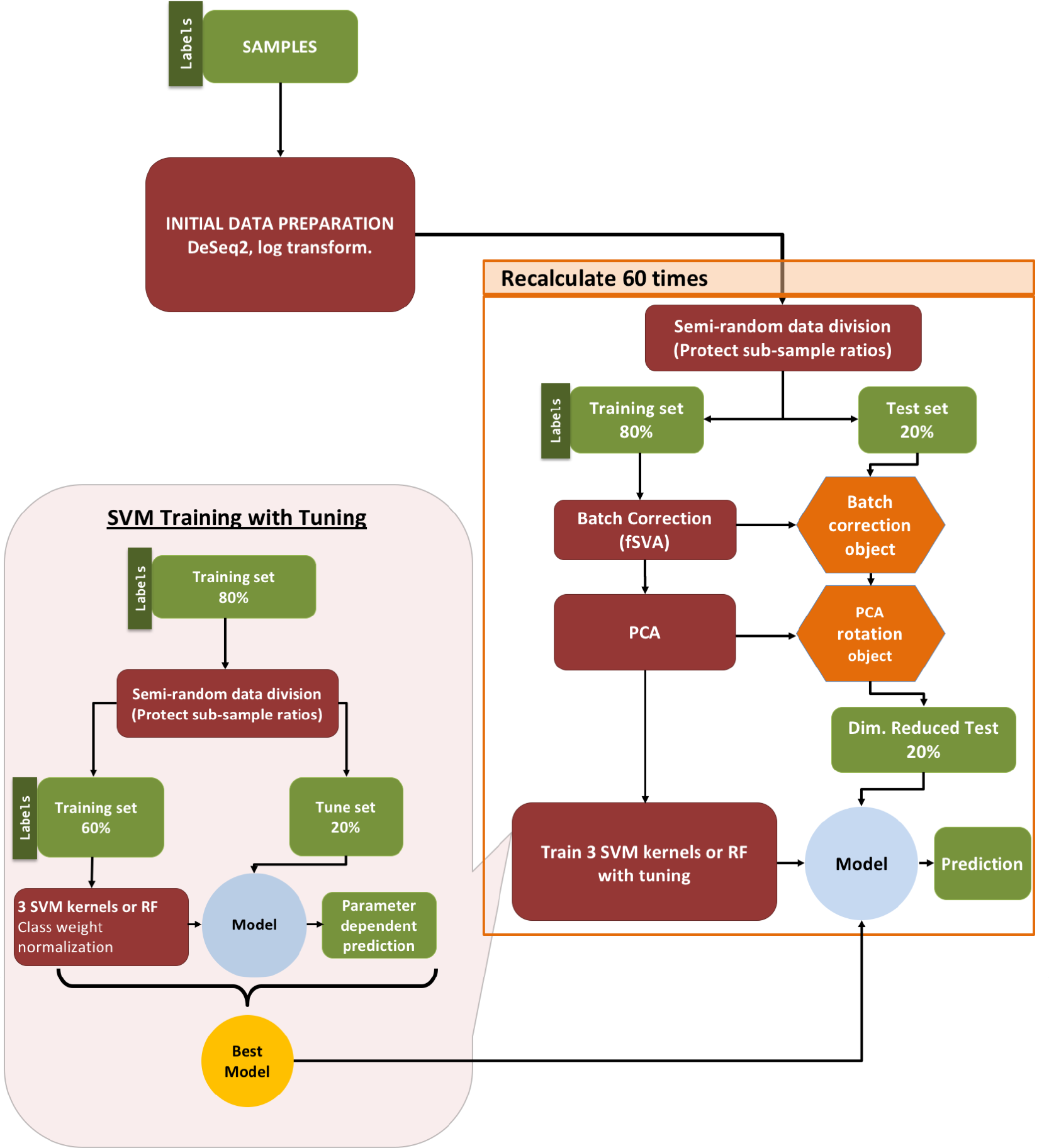
Machine learning pipeline. Our pipeline can be separated into three parts: (i) initial data preparation, (ii) training and prediction, and (iii) model tuning. After (i) initial data preparation, the samples are (ii) semi-randomly (preserving sub-sample ratios) separated into 2 parts, the training & tune set and the test set. After applying fSVA and PCA to the training data, we train supervised SVM or random forest models via tuning. After obtaining the tuned model we make predictions on the test data that has been batch corrected (via fSVA) and rotated (via PCA). This whole process is repeated 60 times to collect statistics on model performance. For model tuning (iii), the training & tune data set is similarly divided semi-randomly into training and tune datasets. The tuning procedure is repeated 10 times and the model that performs best on average during the 10 repeats is considered the winning model and is used for prediction on the test data.

### Growth conditions can be predicted accurately from both mRNA and protein abundances

After constructing our analysis pipeline, we first asked whether there were major differences in the performance of different machine learning approaches. We tested four different machine learning models, three based on Support Vector Machines (SVMs) with different kernels (radial, sigmoidal, and linear) and the fourth using random forest classification. We trained models to predict [7,20] the entire four-dimensional condition vector at once for a given sample, and we used the multi-class macro *F*_1_ score [21] to quantify prediction accuracy. The *F*_1_ score is the harmonic mean of precision and recall. It approaches zero if either quantity approaches zero, and it approaches one if both quantities approach one (representing perfect prediction accuracy). We note that this score is highly conservative as it will classify a prediction as incorrect if a single variable is incorrectly predicted, even if the predictions for the remaining three variables of interest are correct. We assessed model performance during the tuning stage of our pipeline by recording which model had the best *F*_1_ score for each tuning run (S1 and S2 Figs). At the tuning stage, we found that the SVM model with a radial kernel clearly outcompeted the other models when fit to mRNA data, and the random forest model outcompeted the other models when fit to protein data (Table 1).

**Table 1:** Winning-model distributions at the tuning stage. Numbers show the number of times out of 60 independent runs that each given model had the highest *F*_1_ score in the tuning process. Results are shown separately for predictions on the mRNA and the protein data. The ties are counted for all the winner models as a result the sums are bigger than 60

| Model | mRNA | Protein |
| --- | --- | --- |
| SVM, radial kernel | 53 | 8 |
| SVM, sigmoidal kernel | 6 | 41 |
| SVM, linear kernel | 0 | 3 |
| Random Forest | 1 | 13 |

We next compared the *F*_1_ scores for model predictions applied to the test set. When using mRNA abundance data alone, the distribution of *F*_1_ scores from our 60 independent replications were centered around a value of 0.7 (Fig 3). The *F*_1_ score distributions were virtually identical for the three SVM models and were somewhat lower for the random forest model. Model performance on test data using only protein abundance measurements was slightly worse than those achieved with mRNA abundance data. However, it is important to note that the protein abundance data contains fewer conditions overall, which may partially explain the decreased predictive accuracy of the protein-only model—a point to which we return to later.

**Figure 3:**
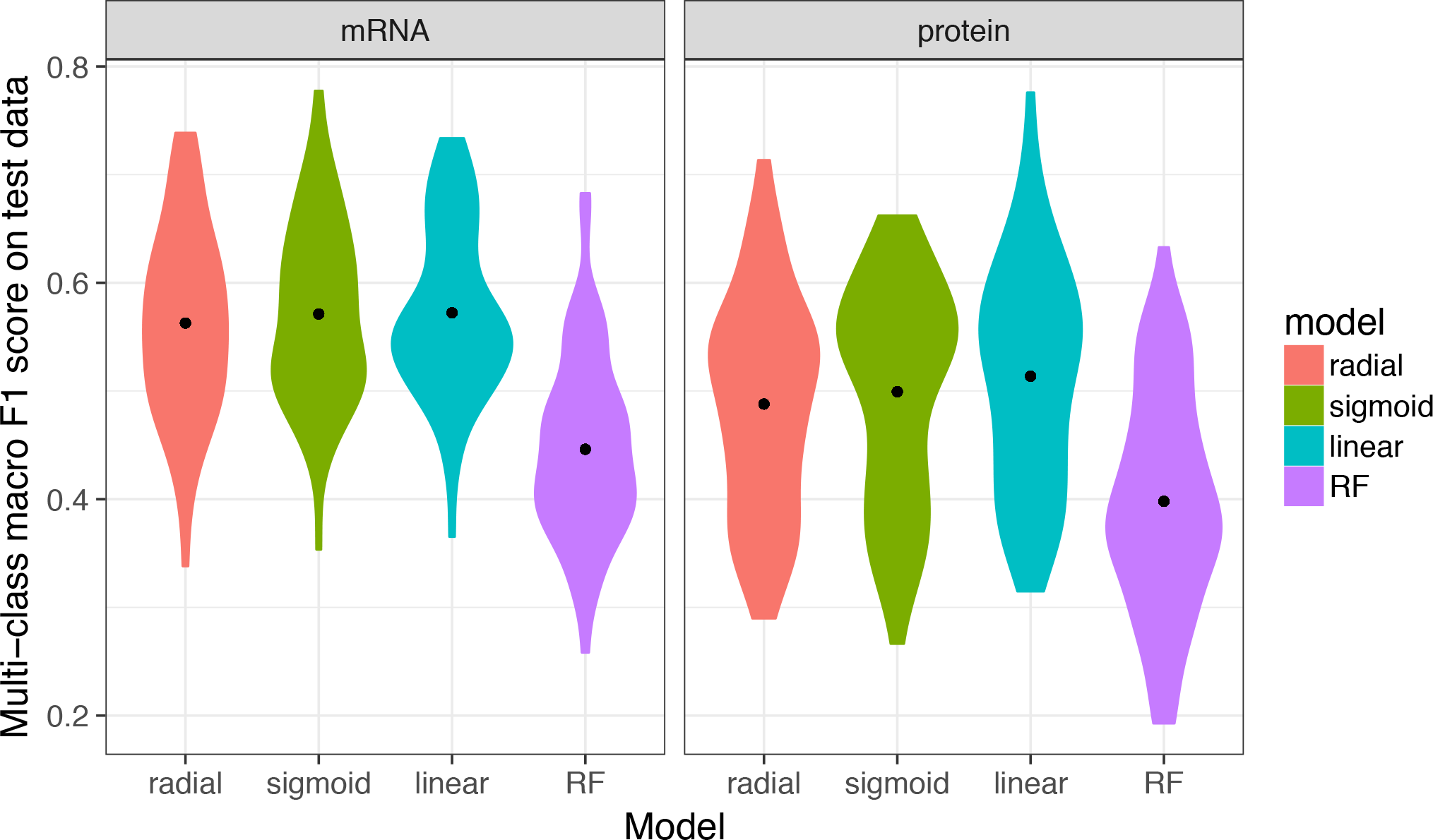
Performance of multi-class predictions. Distributions of multi-class macro *F*_1_ score for prediction of growth conditions from mRNA or protein abundances, using four different machine-learning algorithms (SVM with radial, sigmoidal, or linear kernel, and random forest [RF] models). For each model type, 60 independent models were trained on 60 independent subdivisions of the data into training and test sets. We found that random forest models consistently performed worse than SVM models, and predictions based on mRNA data were slightly better than predictions based on protein data. The black dots represent the mean *F*_1_ scores.

In addition to assessing the overall predictive power using *F*_1_ scores, we also recorded the percentage of times specific growth conditions were accurately or erroneously predicted, and we report these results in the form of a confusion matrix (Fig 4). Here, the column headings at the top show the predicted condition from the model on the test set and the rows show the true experimental condition. The numbers and shading in the interior of the matrix represent the percentage of cases that a given experimental condition was predicted to be a certain growth condition. The numbers within each row add up to 100. The large numbers/dark colorings along the diagonal highlight the high percentage of true positive predictions whereas any off-diagonal elements represent incorrect predictions. We found that the erroneous off-diagonal predictions are partially driven by the uneven sampling of different conditions in the original dataset. Even though we used sample-number-adjusted class weights in all fitted models, we observed a trend of increasing fractions of correct predictions with increasing number of samples available under training (S3 Fig).

**Figure 4:**
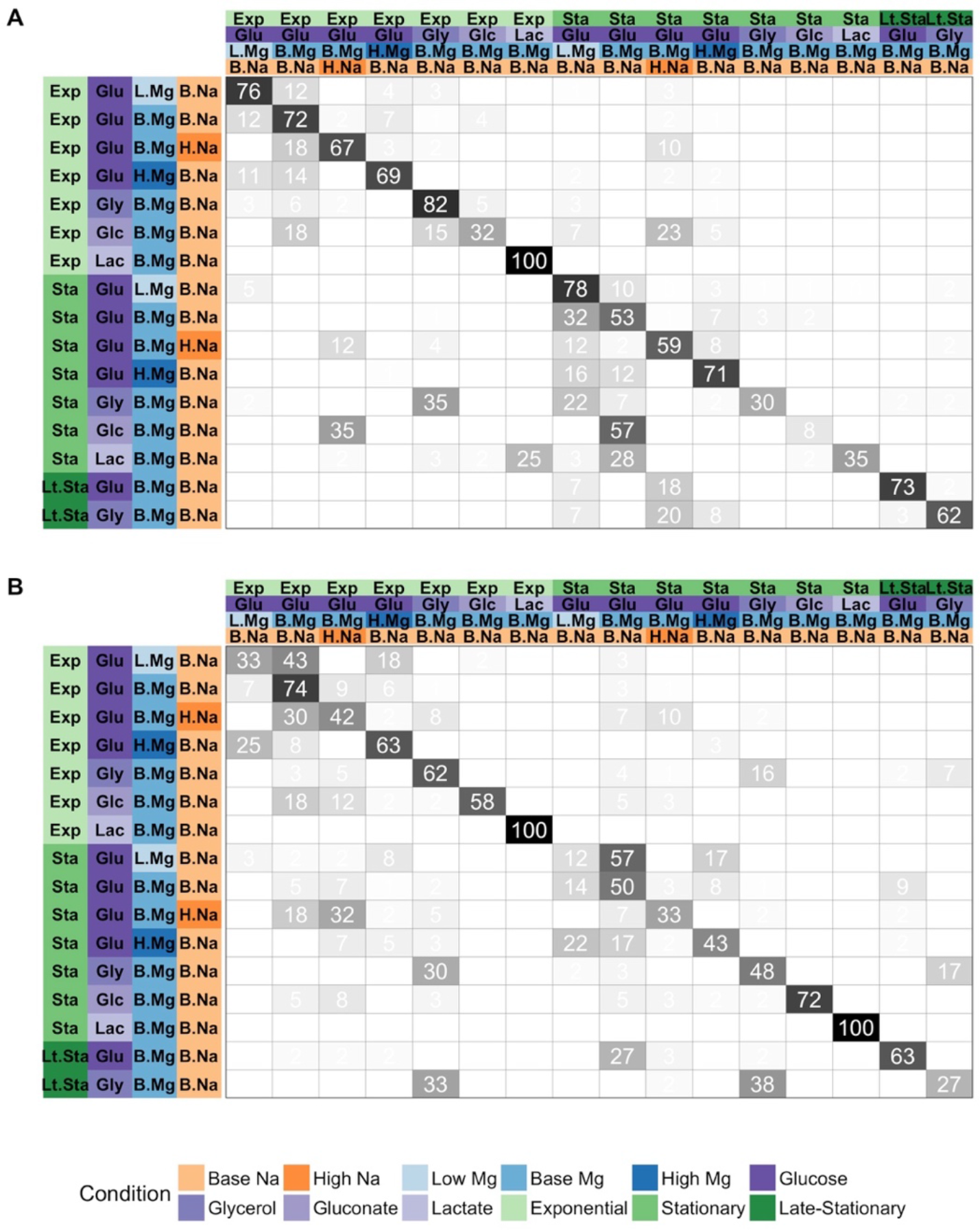
Prediction accuracy for specific growth conditions. In each matrix, rows represent true conditions and columns represent predicted conditions. The numbers in the cells and the shading of the cells represent the percentage (out of 60 independent replicates) with which a given true condition is predicted as a certain predicted condition. (A) Predictions based on mRNA abundances. Results are shown for the SVM with radial kernel, which was the best performing model in the tuning process on mRNA data, where it won 55 of 60 independent runs. In this sub-figure, the average of the diagonal line is 60.5% and corresponding multi-class macro *F*_1_ score is 0.61. (B) Predictions based on protein abundances. Results are shown for the SVM with sigmoidal kernel, which was the best performing model in the tuning process on protein data, where it won 41 of 60 independent runs. In this sub-figure, the average of the diagonal line is 55.1% and corresponding multi-class macro *F*_1_ score is 0.56.

As we previously noted, the *F*_1_ score quantifies accuracy by only considering perfect predictions (i.e. when all 4 features are correctly predicted). A sample that is incorrectly classified for all four factors is thus treated the same as one that only differs from the true set of features by a single incorrect factor. In practice, we observed that the majority of incorrect predictions differed from their true condition vector by only a single value (S4 Fig).

### Joint consideration of mRNA and protein abundances improves model accuracy

We next asked whether predictions could be improved by simultaneously considering mRNA and protein abundances. To address this question, we limited our analysis to the subset of 102 samples for which both mRNA and protein abundances were available, and ran our analysis pipeline for mRNA abundances only, protein abundances only, and for the combined dataset containing both mRNA and protein abundances. For all four machine-learning algorithms, protein abundances yielded significantly better predictions than mRNA abundances (Fig 5, Table 2). This is in contrast to Fig 3, where we saw increased accuracy using mRNA abundance data. However, as previously noted, our dataset contains a larger number mRNA abundance samples, which results in a larger amount of training data. When compared on the same exact conditions—as depicted in Fig 5—protein abundance data appears to be more valuable for discriminating between different growth conditions. Notably, the combined dataset consisting of both mRNA and protein abundance measurements yielded the best overall predictive accuracy, irrespective of machine-learning algorithm used (Fig 5, Table 2).

**Figure 5:**
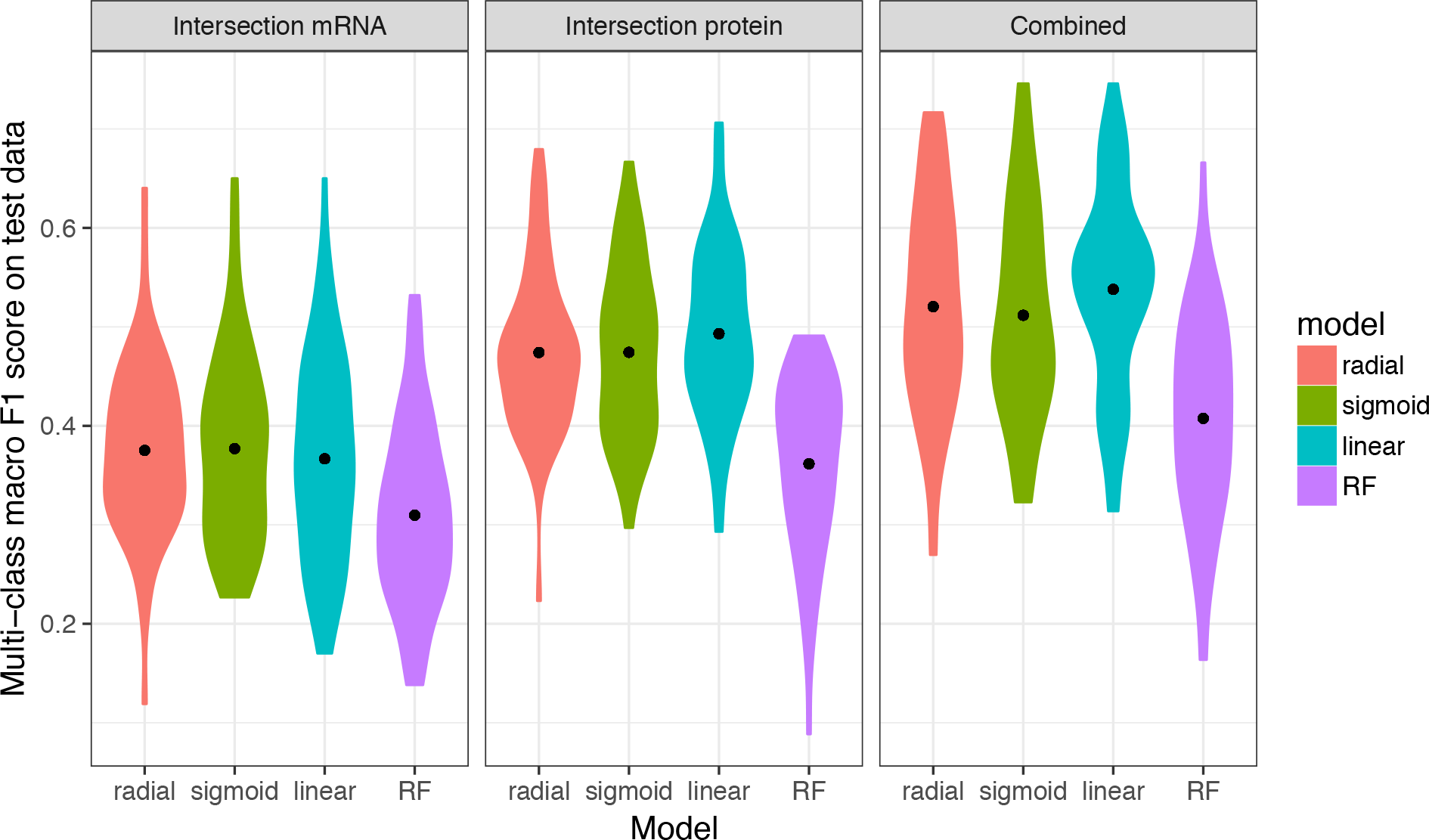
Models trained on both mRNA and protein data perform better than models trained on only one data type. The 102 samples for which we have both protein and mRNA abundances were used to compare the performance of machine learning models based on only mRNA, only protein, and mRNA and protein data combined (left to right, respectively). Regardless of the machine learning model used, prediction performance was higher for models that use protein data compared to mRNA data. Further, using both mRNA or protein data resulted in higher predictive power compared to either alone. Statistical significance of these differences is reported in Table 2.

**Table 2:** Statistical significance of comparisons shown in Figure 5. Distributions of multi-class macro *F*_1_ scores were compared using t-tests. The adjusted *P* value reports the false discovery rate (FDR). All comparisons are statistically significant after correction for multiple testing via FDR.

| Model | Comparison | $P$ value | Adjusted $P$ value |
| --- | --- | --- | --- |
| SVM, radial kernel | mRNA vs protein | 1.943E-09 | 4.663E-09 |
| SVM, radial kernel | mRNA + protein vs mRNA | 3.908E-13 | 2.345E-12 |
| SVM, radial kernel | mRNA + protein vs protein | 8.425E-03 | 1.087E-02 |
|  |  | 3.327E-08 | 6.654E-08 |
| SVM, sigmoidal kernel | mRNA vs protein |  |  |
| SVM, sigmoidal kernel | mRNA + protein vs mRNA | 3.088E-11 | 1.235E-10 |
| SVM, sigmoidal kernel | mRNA + protein vs protein | 3.517E-02 | 3.517E-02 |
|  |  | 4.728E-11 | 1.418E-10 |
| SVM, linear kernel | mRNA vs protein |  |  |
| SVM, linear kernel | mRNA + protein vs mRNA | 1.595E-15 | 1.914E-14 |
| SVM, linear kernel | mRNA + protein vs protein | 9.441E-03 | 1.087E-02 |
|  |  | 1.818E-03 | 2.727E-03 |
| Random forest | mRNA vs protein |  |  |
| Random forest | mRNA + protein vs mRNA | 1.928E-07 | 3.306E-07 |
| Random forest | mRNA + protein vs protein | 9.968E-03 | 1.087E-02 |

When considering the confusion matrices for the three scenarios (mRNA abundance, protein abundance, and combined), we found that many of the erroneous predictions arising from mRNA abundances alone were not that common when using protein abundances and vice versa (S5 and S6 Figs). For example, when using mRNA abundances, many conditions were erroneously predicted as being exponential phase, glycerol, base Mg^2+^, base Na^+^, or as stationary phase, glucose, base Mg^2+^, high Na^+^; these particular predictions were rare or absent when using protein abundances. By contrast, when using protein abundances, several conditions were erroneously predicted as being stationary phase, glycerol, base Mg^2+^, base Na^+^, and these predictions were virtually absent when using mRNA abundance data. For predictions made from the combined dataset, erroneous predictions unique to either mRNA or protein abundances were generally suppressed, and only those predictions that arose for both mRNA and protein abundances individually remained present in the combined dataset (S7 Fig).

### Prediction accuracy differs between environmental features

We also assessed the sources of inaccuracy in our models. As previously noted, the majority of incorrect predictions differed by only a single factor. The environmental features that accounted for most of these single incorrect predictions were Mg^2+^ concentration for the protein-only data and carbon sources for mRNA-only data. Moreover, growth phase (e.g. exponential, stationary, late-stationary) is not strictly an environmental variable and using this as a feature may partially skew our results if the goal is to predict *strictly external* conditions.

We thus trained and tested separate models using only exponential or only stationary phase datasets and asked to what extent these models could predict the remaining 3 environmental features (carbon source, [Mg^2+^], and [Na^+^]). We found that prediction accuracy was consistently better for models trained on exponential-phase samples compared to models trained on stationary-phase samples, irrespective of the machine-learning algorithm used or the data source (mRNA, protein abundances, or both) (Fig 6). This observation implies that *E. coli* gene expression patterns during stationary phase are less indicative of the external environment compared to cells experiencing exponential growth. A notable caveat is that we have fewer stationary phase samples and this decrease in accuracy may partially be due to the size of the training dataset. Even despite the lower accuracies, however, predictive accuracy from models trained solely on stationary phase cells was still much higher than random expectation, illustrating that quiescent cells retain a unique signature of the external environment for the conditions studied.

**Figure 6:**
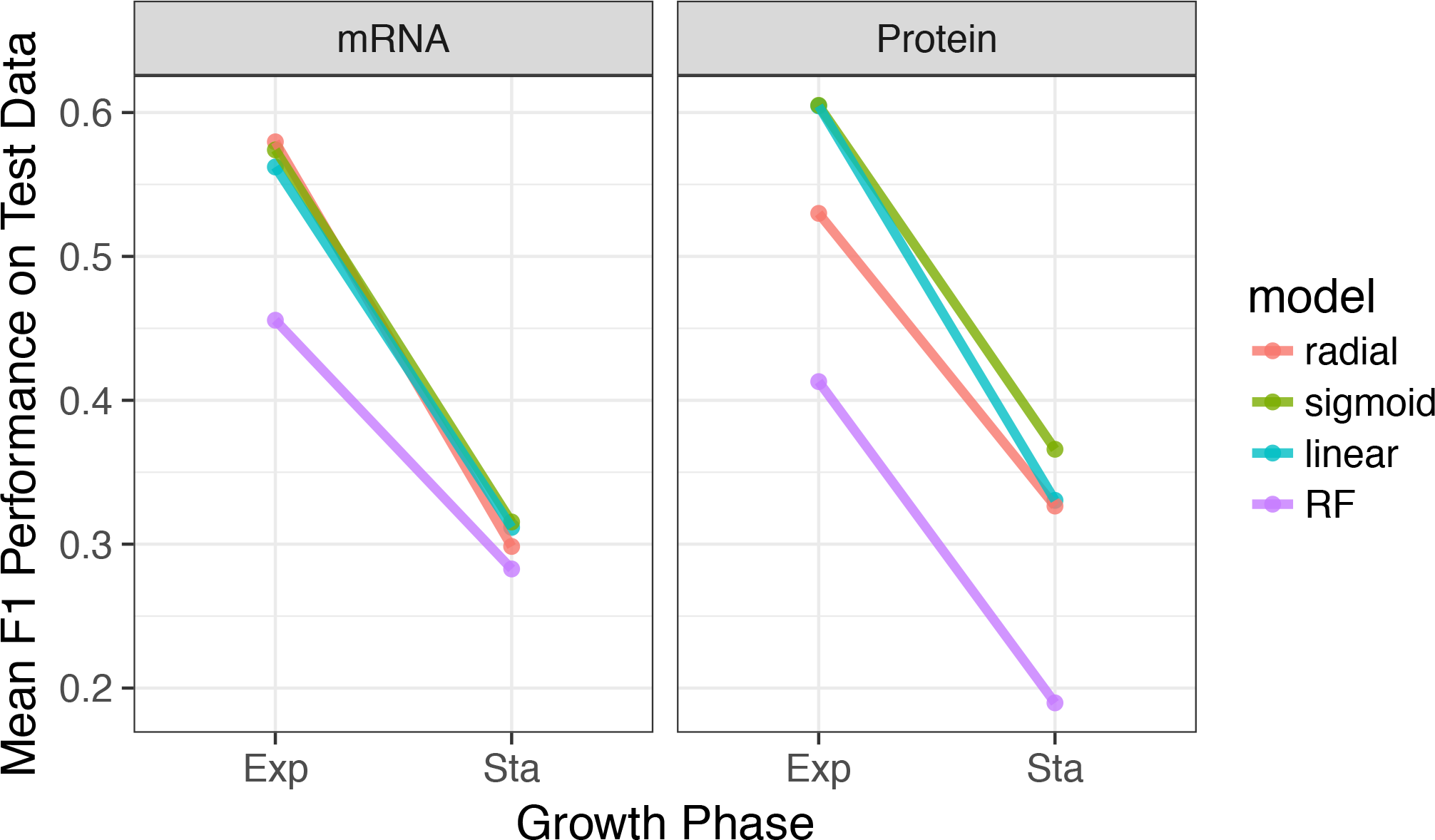
Prediction accuracy systematically declines from exponential to stationary. We separated data by growth phase and then trained models to predict carbon source, magnesium level, and sodium level within each growth phase. Regardless of machine-learning model data source (mRNA or protein), prediction accuracy was substantially lower for stationary-phase samples than for exponential-phase samples. For each model and growth phase, dots show the mean *F*_1_ score over 60 replicates and lines connect mean *F*_1_ scores calculated for the same model.

To better understand which conditions were the most problematic to predict, we constructed models to predict only *individual* features rather than the entire set of 4 features. When making predictions based on mRNA abundances only, models were most accurate in predicting growth phase and least accurate for carbon source, with Mg^2+^ and Na^+^ concentration falling between these two extremes. By contrast, when making predictions based on protein abundances, the most predictable feature was carbon source, the least predictable was Mg^2+^ concentration, and Na^+^ concentration and growth phase fell in-between these two extremes (Fig 7, S8 Fig). Finally, for the combined mRNA and protein abundance dataset, we found that accuracy for carbon source and Mg^2+^ concentration generally fell between the accuracies observed using mRNA and protein abundances individually. By contrast, accuracies for the Na^+^ concentration and growth phase were generally as good as—or better than—the prediction accuracies of the individual datasets (S9 Fig). Together, these findings highlight that mRNA and protein abundances differ in their ability to discriminate between particular environmental conditions.

**Figure 7:**
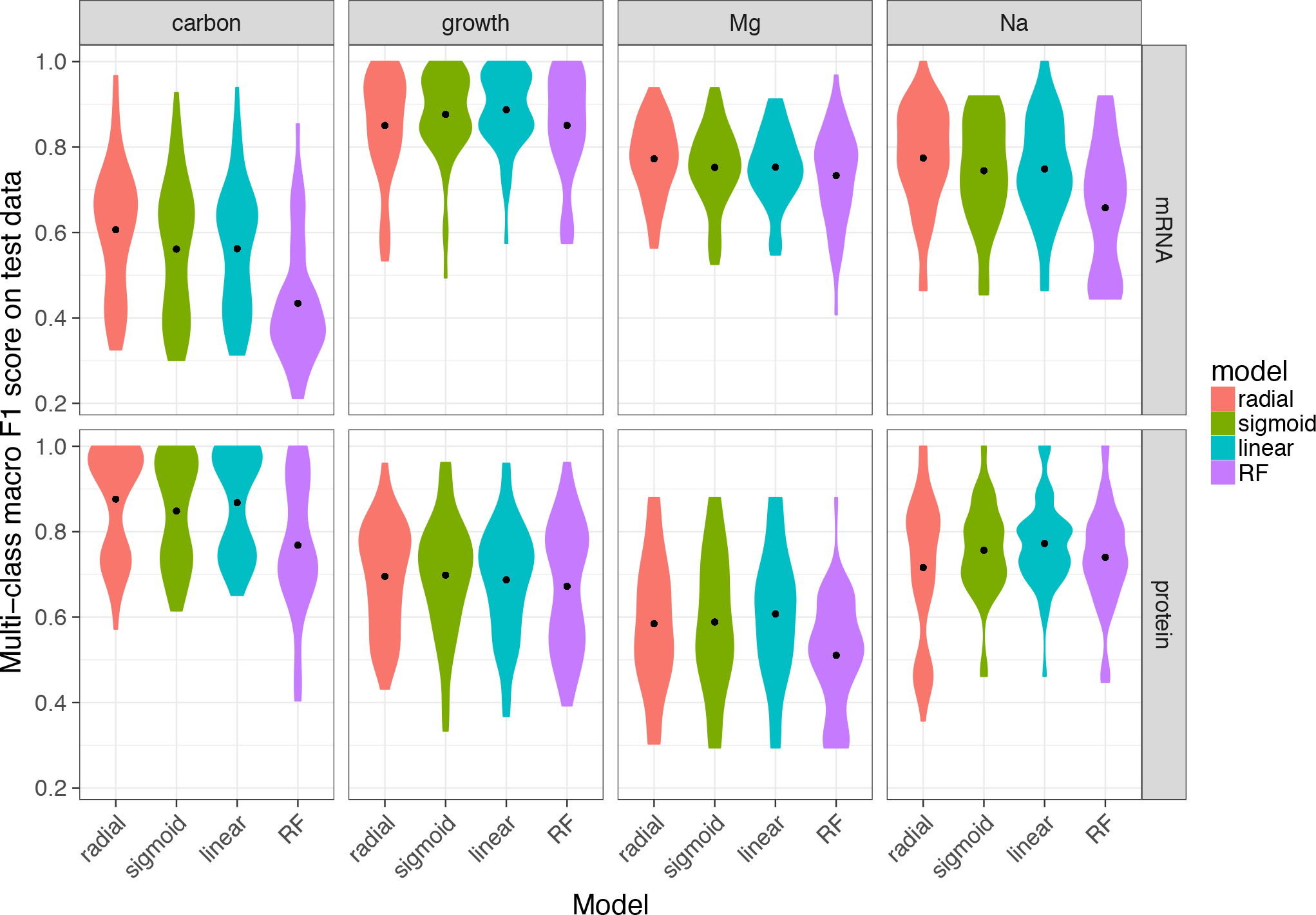
Model performance on univariate predictions. The multi-class macro *F*_1_ score of tuned models over test data for four individual conditions: carbon source, growth phase, Mg^2+^ levels, and Na^+^ levels. To keep mRNA-based and protein-based predictions comparable, we used the 102 samples with both mRNA and protein abundances for this analysis. Note that we used the multi-class macro *F*_1_ score even for univariate predictions, by averaging the component *F*_1_ scores for the individual outcomes, such as the different carbon sources.

### Model validation on external data

The samples that we studied throughout this manuscript are fairly heterogeneous and were collected by different individuals over a span of several months/years. However, different sample types were still analyzed within the same labs, by the same protocols, and thus may be more consistent than one might expect from data collected and analyzed independently by different labs—which would be an ultimate goal of future applications of this methodology. We thus applied our best-fitting protein abundance model to analyze protein data with *similar* conditions that was independently collected and analyzed [7]. Since this external dataset did not contain measurements for all of the 4196 proteins that we measured and constructed our model on, we tested two alternative approaches of applying our model to the external data. For the first approach, we filled the missing parts of the external data with the median values of our in-house data before making predictions. In the second approach, we restricted our training dataset to only include proteins that appeared in the external validation data set. These two approaches lead to comparable results (Fig 8). Notably, our model made mostly correct predictions on this dataset. The model was most accurate at distinguishing between different growth phase data, and moderately accurate at distinguishing Na^+^ concentration and carbon source. The external data did not have variation in Mg^2+^ levels, however, and our model incorrectly predicted several samples to have high Mg^2+^.

**Figure 8:**
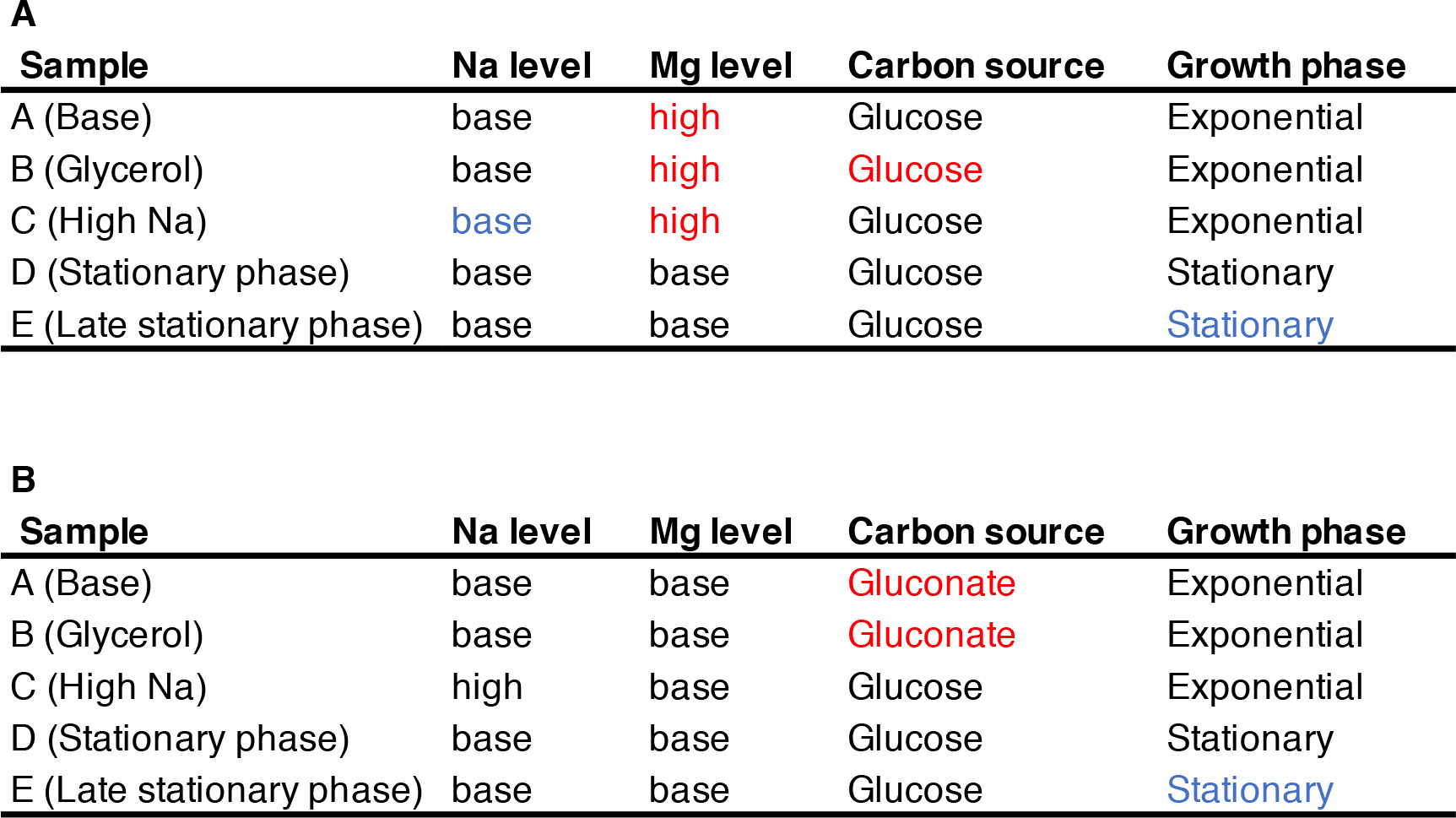
Performance of the protein model on external data. For each of the five external samples we matched to conditions in our dataset, we show the predicted sodium level, magnesium level, carbon source, and growth phase. Black text indicates a correct prediction. Red text indicates an incorrect prediction. Blue text indicates a prediction for a condition where the external data falls between two categories in our data (see Methods for details). (A) Predictions using a model trained on our complete dataset. Any missing protein abundances in the external test data were replaced by the median values from the training dataset. (B) Predictions using a model that was trained on our complete dataset using only the subset of proteins that were present in the external test data.

## Discussion

Our central goal in this manuscript was to determine whether gene expression measurements from a single species of bacterium are sufficient to predict environmental growth conditions. We analyzed a rich dataset of 152 samples for mRNA data and 105 samples for protein data across 16 distinct laboratory conditions as a proof-of-concept. We could show that *E. coli* gene expression is responsive to external conditions in a measurable and consistent way that permits identification of external conditions from gene signatures alone using supervised machine learning techniques. While *E. coli* is a well-characterized species, our analysis relies on none of this *a priori* knowledge. It is thus likely that increasing the number and diversity of training samples and conditions will produce further improvements in accuracy and discrimination between a wider array of conditions.

Interestingly, we found that consideration of mRNA and protein datasets alone are sufficient to produce accurate results, but that joint consideration of both datasets results in superior predictive accuracy. This finding implies that post-transcriptional regulation is at least partially controlled by external conditions, which has been observed by previous studies that have investigated multi-omics datasets [12,20,22,23]. Such regulation may result from post-translational modifications [24], stress coping mechanisms [25], differential translation of mRNAs, or protein-specific degradation patterns.

An important finding that we discovered was that cellular growth phase places limits on the predictability of external conditions, with stationary phase cells being particularly difficult to distinguish from one another irrespective of their external conditions. A possible explanation for this behavior might be associated with endogenous metabolism, whereby stationary phase cells start to metabolize surrounding dead cells instead of the provided carbon source. This new carbon source, which is independent of the externally provided carbon source, may suppress the differences between the cells in different external carbon source environments [26,27]. Another reason for this behavior might be related to strong coupling between gene expression noise and growth rate. Multiple studies have concluded that lower growth rates are associated with higher gene expression noise, which might be a survival strategy in harsh environments [28]. Negative correlations between population average gene expression and noise have been shown for *E. coli* and *Saccharomyces cerevisiae*, lending support for this theory [29,30]. Finally, we note that stationary phase cells have likely depleted the externally supplied carbon sources after several weeks of growth. The similarity of stationary phase cells to other stationary phase cells may be a consequence of them inhabiting more similar chemical environments to one another compared to during exponential growth where nutrient concentrations are more varied across conditions. Nevertheless, discrimination of external environmental factors in stationary phase cells was still much better than random—indicating that these populations continue to retain information about the external environment despite their overall quiescence.

A relevant finding to emerge from our study is that different features of the environment may be more or less easy to discriminate from one another and this discrimination may depend on which molecular species is being interrogated. Growth phase, for instance, can be reliably predicted from mRNA concentrations but similar predictions from protein concentrations were less accurate. A possible explanation for this observation is the fact that mRNAs and proteins have different life-cycles [19,31]. Given the comparably slow degradation rates of proteins, a large portion of the stationary phase proteome is likely to have been transcribed during exponential phase growth. As another example, carbon sources can be reliably predicted from protein concentrations, but the accuracy of carbon source predictions from models trained on mRNA concentrations was more limited. Carbon assimilation is known to be regulated by post-translational regulation [32–34], which may be a possible reason for this finding (Fig 7, S9 Fig).

Despite the fact that we investigated over 150 samples spanning 16 unique conditions, a limitation of our work and conclusions is nevertheless sample size (though our study is comparable to or larger than similar multi-conditional transcriptomic and/or proteomic studies [7,35–37]). The comparison between all of our data with the more limited set that includes only the intersection of samples for which we have both mRNA and protein abundance data (Fig 4 compared to S5 and S6 Figs) indicates that prediction accuracy decreases as the size of our training sets get smaller. This trend indicates that our training set sizes are still ultimately limiting model accuracy. A second possible issue with our study is associated with sample number bias [38–40]. We made corrections with weight factors [41,42] and displayed the multi-class macro *F*_1_ score [43] to account for the fact that some conditions contained more samples, but the predictability of *individual* conditions nevertheless increased with the number of training samples for that particular condition (S3 Fig). This finding again highlights that increasing training data will likely result in higher prediction accuracy.

Our study is a proof-of-principle towards the goal of using gene expression patterns of natural species as a rapid and low-cost method for assessing environmental conditions. Other research has shown that the species repertoire, derived from meta-genomic sequencing, may be useful for determining the presence of particular contaminants [3]. Our findings suggest that further incorporation of species-specific gene expression patterns can likely improve the accuracy of such methods. While genetically engineered strains may play a similar role as environmental biosensors, our study highlights that—with enough training data—the molecular composition of natural populations may provide sufficient information to accurately resolve past and present environmental conditions.

## Materials and Methods

### Data preparation and overall analysis strategy

We used a set of 155 *E. coli* samples previously described [18,19]. Throughout this study, we used different subsets of these samples in different parts of the analysis. For “mRNA only” and “protein only” analyses we used all 152 samples with mRNA abundances and all 105 samples with protein abundances, respectively. For performance comparison of machine learning models between mRNA and protein abundances we used the subset of 102 samples that have both mRNA and protein abundance data. After selecting appropriate subsets of the data for a given analysis, we added abundances from technical replicates, normalized abundances by size factors calculated via DeSeq2 [44], and applied a variance stabilizing transformation [45,46] (VST).

For each separate analysis, we divided the data into two subsets, (i) the training & tune set and (ii) the test set, using an 80:20 split (Fig 2). This division was done semi-randomly, such that our algorithm preserved the ratios of different conditions between the training & tune and the test subsets. We retained the condition labels in the training & tune data (thus our learning was supervised) but we discarded the sample labels for the test set. We then applied frozen Surrogate Variable Analysis [47] (fSVA) to remove batch effects from the samples. This algorithm can correct for batch effects in both the training & tune and the test data, without knowing the labels of the test data. After fSVA, we used principal component analysis [48] (PCA) to define the principal axes of the training & tune set and then rotated the test data set with respect to these axes. We then picked the top 10 most significant axes in the training & tune dataset for learning and prediction. Finally, we trained and tuned our candidate machine learning algorithms with the dimension reduced training & tune dataset and then applied those trained and tuned algorithms on the dimension-reduced test dataset to make predictions. This entire procedure was repeated 60 times for each separate analysis (Fig 2).

We used four different machine learning algorithms: SVM models with (i) linear, (ii) radial, and (iii) sigmoidal kernels, and (iv) random forest models. We used the R package e1071 [49] for implementing SVM models and the R package randomForest [50] for implementing random forest models. SVMs with radial and sigmoidal kernels were set to use the c-classification [51] algorithm.

### Model scoring

Our goal throughout this work was to predict multiple parameters (i.e., growth phase, carbon source, Mg^2+^ concentration, or Na^+^ concentration) of each growth condition at once. Therefore, we could not measure model performance via ROC or precision-recall curves, which assume a simple binary (true/false) prediction. Instead, we assessed prediction accuracy via *F*_1_ scores, which jointly assess precision and recall. In particular, for predictions of multiple conditions at once, we scored prediction accuracy via the multi-class macro *F*_1_ score [21,43,52] that normalizes individual *F*_1_ scores over individual conditions, i.e., it gives each condition equal weight instead of each sample. There are two different macro *F*1 score calculation that have been proposed in the literature. First, we can average individual *F*_1_ scores over all conditions *i* [43]:

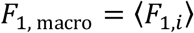

where 〈…〉 indicates the average and the individual *F*_1_ scores are defined as:

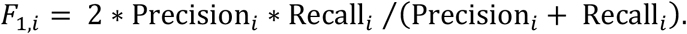

Alternatively, we can average precision and recall and then combine those averages into an *F*_1_ score [21]:

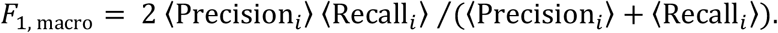

Between these two options, we implemented the first, because it is not clear that individually averaging precision and recall before combining them into *F*_1_ appropriately balances prediction accuracies from different conditions with very different prediction accuracies.

### Model training and tuning

For training, we first divided the training & tune data further into separate training and tuning datasets, using a 75:25 split (Fig 2). As before for the subdivision between training and test data, we did this again semi-randomly, trying to preserve the ratios of individual conditions. We repeated this procedure 10 times to generate 10 independent pairs of training and tuning datasets. Next, we generated a parameter grid for the tuning process. We optimized the “cost” parameter for all three SVM models and the “gamma” parameter for the SVM models with radial and sigmoidal kernels (S1 Fig). For the random forest algorithm, we optimized three parameters; “mtry”, “ntrees”, and “nodesize”.

We trained each of the four machine learning models on all 10 training datasets and made predictions on the 10 tuning datasets. We applied a class weight normalization during training, where class weights are inversely proportional to the corresponding number of training samples and calculated independently for each training run. We calculated macro *F*_1_ scores for each model parameter setting for each tuning dataset and then averaged the scores over all tuning datasets to obtain an average performance score for each algorithm and for each parameter combination. The parameter combination with the highest average *F*_1_ score was considered the winning parameter combination and was subsequently used for prediction on the test dataset (Fig 2).

### Model validation on external data

We validated our predictions against independently published external data [7]. This external dataset consisted of 22 conditions, of which we could match five to our conditions. For all five samples, Mg^2+^ levels were held constant and approximately matched our base Mg^2+^ levels. The first sample used glucose as carbon source, did not experience any osmotic stress (no elevated sodium), and was collected in the exponential growth phase. The second sample used glycerol as carbon source, did not experience any osmotic stress (no elevated sodium), and was collected in the exponential growth phase. The third sample included 50mM sodium, glucose as carbon source, and was collected in the exponential growth phase. Because our high-sodium samples all included 100mM of sodium or more [18], this third sample fell in-between what we consider base sodium and high sodium. Samples four and five used glucose as carbon source, did not experience osmotic stress, and were measured after 24 and 72 hours of growth, respectively. In our samples, we defined stationary phase as 24-48 hours and late stationary phase as 1 to 2 weeks [18]. Thus, sample four matched our stationary phase samples and sample five fell in-between our stationary and late-stationary phase samples.

### Statistical analysis and data availability

All statistical analyses were performed in R. All processed data and analysis scripts are available on GitHub: https://github.com/umutcaglar/ecolimultiplegrowthconditions (permanent archived version available via zenodo: 10.5281/zenodo.1294110). mRNA and protein abundances have been previously published [18,19]. Raw Illumina read data and processed files of read counts per gene are available from the NCBI GEO database [53] (accession numbers GSE67402 and GSE94117). Mass spectrometry proteomics data are available via PRIDE [54] (accession numbers PXD002140 and PXD005721).

## Acknowledgements

The authors acknowledge support from the Texas Advanced Computing Center (TACC) at The University of Texas at Austin for providing high-performance computing resources.

## Supporting information

**S1 Fig.** Tuning results for predictions based on mRNA data, generated from one of 60 independent runs and chosen for demonstration purposes. Model performance is measured as the mean *F*_1_ score over 10 independent tuning runs. Higher numbers indicate better performance. (A) Tuning results for SVMs with linear kernel. Only the cost parameter was tuned. (B) Tuning results for SVMs with radial kernel. The cost and gamma parameters were tuned. The red dot indicates the winning parameter combination. (C) Tuning results for SVMs with sigmoidal kernel. The cost and gamma parameters were tuned. The red dot indicates the winning parameter combination. (D) Tuning results for random forest models. The mtry, nodesize, and ntrees parameters were tuned. We used three values for ntrees, 1000, 5000, and 10000, shown as three separate panels. The red dot indicates the winning parameter combination.

**S2 Fig.** Tuning results for predictions based on protein data, generated from one of 60 independent runs and chosen for demonstration purposes. (A) Tuning results for SVMs with linear kernel. Only the cost parameter was tuned. (B) Tuning results for SVMs with radial kernel. The cost and gamma parameters were tuned. The red dots indicate the winning parameter combinations. (C) Tuning results for SVMs with sigmoidal kernel. The cost and gamma parameters were tuned. The red dot indicates the winning parameter combination. (D) Tuning results for random forest models. The mtry, nodesize, and ntrees parameters were tuned. We used three values for ntrees, 1000, 5000, and 10000, shown as three separate panels. The red dot indicates the winning parameter combination.

**S3 Fig.** Percentage of correct predictions as a function of the number of samples during training. (A) Predictions based on mRNA abundances. (B) Predictions based on protein abundances.

**S4 Fig.** The error count distribution for mRNA (A) and protein (B) confusion matrices. The number of mis-predicted labels (x-axis) indicates how many of the 4 possible condition variables that an individual prediction got wrong. 0 mis-predicted labels (the majority in both cases) means that model predictions were 100% accurate. In both cases (mRNA and protein), when an incorrect prediction was made, it was most frequently due to a single variable being incorrectly predicted (number of mis-predicted labels with a value of 1) as compared to errors predicting more than one variable for a given condition (2 and 3 mis-predicted labels).

**S5 Fig.** Prediction accuracy for specific growth conditions for intersection mRNA data. Rows represent true conditions and columns represent predicted conditions. The numbers in the cells and the shading of the cells represent the percentage (out of 60 independent replicates) with which a given true condition is predicted as a certain predicted condition. Predictions based on mRNA abundances, generated by using subset of mRNA samples which has matching protein pairs. Results are shown for the SVM with radial kernel, which was the best performing model in the tuning process on mRNA data, where it won 48 of 60 independent runs. In this figure average of the diagonal line is 44.1% and multi class macro *F*_1_ score is 0.43.

**S6 Fig.** Prediction accuracy for specific growth conditions for intersection protein data. Rows represent true conditions and columns represent predicted conditions. The numbers in the cells and the shading of the cells represent the percentage (out of 60 independent replicates) with which a given true condition is predicted as a certain predicted condition. Predictions based on protein abundances, generated by using subset of protein samples which has matching mRNA pairs. Results are shown for the SVM with sigmoid kernel, which was the best performing model in the tuning process on mRNA data, where it won 47 of 60 independent runs. In this figure average of the diagonal line is 52.3% and corresponding multi class macro *F*_1_ score is 0.53.

**S7 Fig.** Prediction accuracy for specific growth conditions for intersection mRNA & protein data. Rows represent true conditions and columns represent predicted conditions. The numbers in the cells and the shading of the cells represent the percentage (out of 60 independent replicates) with which a given true condition is predicted as a certain predicted condition. Predictions based on protein abundances, generated by using subset of mRNA & protein samples which has matching pairs. Results are shown for the SVM with sigmoid kernel, which was the best performing model in the tuning process on combined intersection data, where it won 27 of 60 independent runs. In this figure average of the diagonal line is 56.1% and corresponding multi class macro *F*_1_ score is 0.57.

**S8 Fig.** Prediction accuracy for univariate predictions using intersection mRNA and intersection protein data, as in the main text Figure 7. (A) Prediction of carbon source from mRNA abundances. (B) Prediction of carbon source from protein abundances. (C) Prediction of growth phase from mRNA abundances. (D) Prediction of growth phase from protein abundances. (E) Prediction of Mg^2+^ levels from mRNA abundances. (F) Prediction of Mg^2+^ levels from protein abundances. (G) Prediction of Na^+^ levels from mRNA abundances. (H) Prediction of Na^+^ levels from protein abundances.

**S9 Fig.** Prediction accuracy for univariate predictions based on intersection mRNA abundances, intersection protein abundances, or the combined dataset including both mRNA and protein abundances. Protein abundances are more predictive for carbon source and Mg^2+^ levels, and mRNA abundances are more predictive for Na^+^ levels and growth phase.

